# Repurposing Tranexamic Acid as an Anticancer Agent

**DOI:** 10.1101/2021.10.17.464714

**Authors:** Mary E. Law, Bradley J. Davis, Amanda F. Ghilardi, Elham Yaaghubi, Zaafir M. Dulloo, Mengxiong Wang, Olga Guryanova, Coy D. Heldermon, Ronald K. Castellano, Brian K. Law

**Author notes:** Corresponding Author **Address of Corresponding Author** Brian K. Law, 1200 Newell Drive, Room R5-210 Academic Research Building, University of Florida, Gainesville, FL 32610.

## Abstract

Tranexamic Acid (TA) is a clinically used antifibrinolytic that acts as a lysine mimetic to block binding of Plasminogen with Plasminogen activators, preventing conversion of Plasminogen to its proteolytically activated form, Plasmin. Previous studies suggested that TA may exhibit anticancer activity by blockade of extracellular Plasmin formation. Plasmin-mediated cleavage of the CDCP1 protein may increase its oncogenic functions through several downstream pathways. Results presented herein demonstrate that TA blocks Plasmin-mediated excision of the extracellular domain of the oncoprotein CDCP1. *In vitro* studies indicate that TA reduces the viability of a broad array of human and murine cancer cell lines, and breast tumor growth studies demonstrate that TA reduces cancer growth *in vivo*. Based on the ability of TA to mimic lysine and arginine, we hypothesized that TA may perturb multiple processes that involve Lys/Arg-rich protein sequences, and that TA may alter intracellular signaling pathways in addition to blocking extracellular Plasmin production. Indeed, TA-mediated suppression of tumor cell viability is associated with multiple biochemical actions, including inhibition of protein synthesis, reduced activating phosphorylation of STAT3 and S6K1, decreased expression of the MYC oncoprotein, and suppression of Lys acetylation. These findings suggest that TA or TA analogs may serve as lead compounds and inspire the production of new classes of anticancer agents that function by mimicking Lys and Arg.

## INTRODUCTION

Repurposing FDA-approved drugs as anticancer agents represents a powerful method for rapidly advancing new cancer therapeutics into the clinic especially since many of the safety issues are already established. This is particularly true for a drug like Tranexamic Acid (TA), which is on The WHO List of Essential Medicines and has an excellent safety record while being widely used for over 50 years in a broad range of formulations and dosages in numerous different indications.Tranexamic Acid (TA) is clinically used to suppress hemorrhaging by blocking Plasminogen conversion to Plasmin. TA functions as a lysine side chain mimetic and prevents Plasmin formation by occupying lysine binding sites on Plasminogen (Iwamoto 1975). Due to its ability to mimic the side chains of lysine, and perhaps arginine, yet not being incorporated into protein due to its structure, TA has the potential to alter many biological processes. For instance, basic amino acid residues are present in the nuclear import signals of proteins (Laskey, et al. 1996) and in the consensus recognition sites of many protein kinases, most noteworthy, the AGC family of serine/threonine kinases (Zhang, et al. 2002). Additionally, lysine side chains in proteins are subject to multiple posttranslational modifications that include acetylation, methylation, ubiquitination, and SUMOylation. The “histone code” of epigenetic regulation involves lysine and arginine modifications (Blee, et al. 2015, Izzo and Schneider 2010, Turner 2000, Tweedie-Cullen, et al. 2012). Given the important role of lysine and its modifications in the control of multiple cellular processes, we hypothesized that in addition to its widespread use to suppress bleeding, TA may inhibit the enzymes that carry out lysine modification in proteins in a competitive manner and suppress the function of lysine/arginine-rich protein-protein recognition motifs.

While TA has been used to manage cancer and cancer treatment-associated sequela in patients (Jaffer, et al. 2021, Kikuchi, et al. 1986, Lohani, et al. 2020, Lohani, et al. 2021, Longo, et al. 2018, Oertli, et al. 1994, Soma, et al. 1980, Wu, et al. 2006), it has never been used clinically as an anti-cancer therapeutic, and considerations of TA anticancer actions have focused on its blockade of Plasmin production. The goals of the studies carried out herein were three-fold: 1. To examine the impact of TA treatment on the viability of cancer cells to evaluate the possibility of repurposing TA as an anticancer agent, 2. To determine if TA may provide a lead compound to serve as the basis of derivatives with greater anticancer efficacy and potency, and enhanced selectivity for specific downstream responses, and 3. Because of its unusual mechanism of action, test whether TA may prove useful in combination anticancer regimens. Given our previous work with a class of anticancer compounds termed Disulfide bond Disrupting Agents (DDAs) (Besse, et al. 2019, Ferreira, et al. 2015, Ferreira, et al. 2017, Wang, et al. 2019b, Wang, et al. 2019c) and observations below that TA and DDAs have partially overlapping biochemical mechanisms of action, we evaluated the anticancer of TA and DDA mono-and combination therapies in an animal model of breast cancer. The results obtained show that TA suppresses the viability of a broad array of human and cancer cell lines, exhibits previously unreported effects on cell signaling pathways that control cancer cell proliferation and survival, and modulates acetylation status of lysine residues in a subset of acetylated proteins. TA and the DDAs tcyDTDO and dMtcyDTDO strongly suppress the growth of breast tumors individually and may be useful on combination.

## MATERIALS AND METHODS

### MTT Cell viability assays

MTT cell viability assays were performed by plating cell lines at 7,500 cell/well in 96-well plates, followed by the treatment of cells with increasing concentrations of TA for 72 h at 37°C. After removal of the cell media, cells were incubated with 0.5 mg/mL MTT (3-(4,5-dimethylthiazol-2-yl)-2,5-diphenyltetrazolium bromide) (Biomatik, Wilmington, DE USA) in PBS for 3 h. The MTT solution was subsequently removed and the MTT formazan product was dissolved in 100 μL of DMSO. Absorbance (570 and 690 nm) of the MTT formazan product was assessed in a plate reader.

### *In vivo* tumor studies

012/LVM2/LR10 tumors were generated by injection of 1□×□10^6^ 012/LVM2/LR10 cells into the #4 mammary fat pads of NOD-SCID-gamma (NSG) mice from Jackson Laboratories (Bar Harbor, ME USA), as described previously (Wang, et al. 2019b). In the tumor study in Fig. 1F, G, mice bearing HCI-001/LVM2/LR10 breast tumors were treated once daily with 375 mg/kg TA by oral gavage for 12 days followed by breast tumor removal. In the tumor study in Fig. 4A, tumors were generated using the 012/LVM2/LR10 cancer cell model as above and the animals were treated with vehicle (peanut oil), 375 mg/kg TA, 10 mg/kg dMtcyDTDO, or 375 mg/kg TA + 10 mg/kg dMtcyDTDO once daily by oral gavage. Tumor dimensions were recorded daily at the time of treatment.

**Fig. 1:**
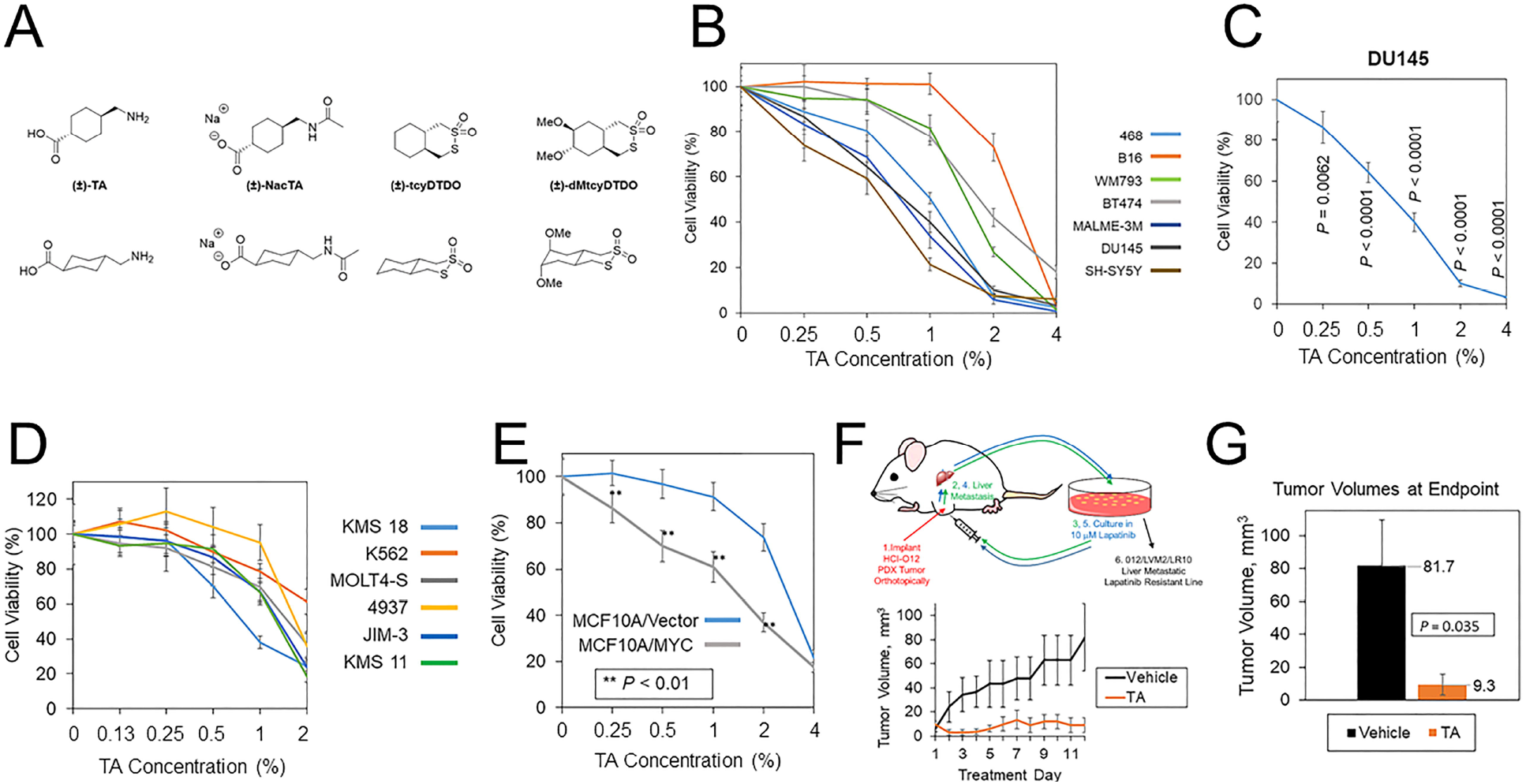
TA decreases cancer cell viability and breast tumor growth. A. Chemical structures of TA, *N*-acetylated TA (NacTA), and the DDAs tcyDTDO and dMtcyDTDO. B. The indicated panel of cancer cell lines were treated for 72 h with different TA concentrations and cell viability was measured using the MTT assay. C. Viability assay on DU145 cells performed as in A. examining the statistical significance of TA induced reduction in cell viability. Statistically significant reductions in viability were observed at all TA concentrations tested, down to 0.25% w/v. D. Effects of TA on the viability of a panel of leukemia, lymphoma, and multiple myeloma cell lines. E. Cell viability assay showing that TA more potently reduces the viability of MCF10A mammary epithelial cells overexpressing MYC compared with the vector control line. Statistically significant differences between the lines were observed at 0.25, 0.5, 1, and 2% TA. F. Tumor growth study in which mice bearing orthotopic 012/LVM2/LR10 mammary tumors were treated once daily with 375 mg/kg TA. The upper panel illustrates how the 012/LVM2/LR10 cancer model was derived. Tumor dimensions were measured daily and tumor volumes calculated. Averages of within-group tumor volumes are presented and error bars represent standard error of the mean. G. Tumor volumes measured at day 12 exhibit a statistically significant TA-mediated reduction in tumor volume (*P* = 0.035), with an approximately nine-fold smaller tumor volume in the TA group than the vehicle group.

### Protein Synthesis Assays

Protein synthesis assays were performed by measurement of the incorporation of ^3^H-Leucine (Perkin Elmer, Waltham, MA USA) into protein, as detailed previously (Law, et al. 2000a, Law, et al. 2000b).

### Thymidine Incorporation Assays

Thymidine incorporation assays are described in a previous publication (Corsino, et al. 2009).

### P300 Acetylation Assays

P300 Acetylation assays were performed by incubating the reaction buffer (50 mM Tris, pH 8.0, 10% glycerol, 0.1 mM EDTA, and 1 mM DTT), 0.5 μg p300 (ENZO Life Sciences, Farmingdale, NY USA), 0.5 μg Histone H3 (NEB, Ipswich, MA USA), and 60 μM Acetyl CoA (Sigma-Aldrich, St. Louis, MO USA) in the presence or absence of TA (Chem-Impex International, Inc., Wood Dale, IL USA), NacTA, or tcyDTDO for 2 h at 30°C. Reactions were terminated by the addition of 2X SDS sample buffer and boiling for 10 min, followed by immunoblot analysis employing a 45 min transfer to 0.2 μm nitrocellulose using CAPS buffer, pH 11.

### NacTA Synthesis

#### General methods

Reagents and solvents were purchased from commercial sources without further purification unless otherwise specified. ^1^H and ^13^C NMR spectra were recorded using commercially obtained D_2_O (Cambridge Isotope Laboratories, Inc.) on a Bruker 600 spectrometer (^1^H at 600 MHz) and Bruker 400 spectrometer (^13^C at 101 MHz), respectively. Chemical shifts (*δ*) are given in parts per million (ppm) relative to TMS and referenced to residual protonated D_2_O (*δ*H = 4.79 ppm).

Coupling constants (*J*) are given in Hz; spin multiplicities are presented by the following symbols: s (singlet), bs (broad singlet), d (doublet), t (triplet), q (quartet), and m (multiplet). Electrospray ionization (ESI) high resolution mass spectra (HRMS) were recorded on an Agilent 6200 ESI-TOF instrument operating in positive or negative ion mode as stated.

#### Synthesis of sodium (1*r*,4*r*)-4-(acetamidomethyl)cyclohexane-1-carboxylate, NacTA

The preparation of NacTA was adapted from a procedure reported by Kusakabe, *et al* (Kusakabe 2010). Tranexamic acid (1.90 g, 12.1 mmol) was suspended in acetic anhydride (9.70 mL, 103 mmol) and concentrated H_2_SO_4_ (5.00 μL) was carefully added. The suspension was stirred for 18 h at rt. Water (10.0 mL) was added and the mixture was stirred for 1 h at rt to decompose any remaining acetic anhydride. The solution was concentrated under vacuum, and the residue was further dried with the addition of toluene and subsequent removal of solvent under reduced pressure. The resulting precipitate was collected by vacuum filtration and washed with ethyl ether to give NacTA as a white powder. For the sodium salt preparation, NacTA was suspended in water and an aqueous solution of 2 M NaOH was added with stirring at rt until the pH increased to 13. The suspension was then stirred for an additional 1 h at rt until the entire solid dissolved. The solvent was then evaporated under reduced pressure and the precipitate was recrystallized in ethanol to remove remaining NaOH salt.The white precipitate obtained was then collected by vacuum filtration and washed with ethanol to afford the product as a white solid. (1.30 g, 5.80 mmol, 48% yield). ^1^H NMR (600 MHz, D_2_O) *δ* 3.01 (d, *J* = 6.8 Hz, 2H), 2.08 (tt, *J* = 12.2, 3.5 Hz, 1H), 1.98 (s, 3H), 1.90–1.84 (m, 2H), 1.77 (dd, *J* = 13.2, 3.6 Hz, 2H), 1.52–1.42 (m, 1H), 1.31 (qd, *J* = 13.0, 3.7 Hz, 2H), 0.96 (qd, *J* = 12.8, 3.6 Hz, 2H). The NH proton was not observed. ^13^C NMR (101 MHz, D_2_O) *δ* 186.50, 173.97, 46.92, 45.53, 36.78, 29.49, 29.23, 21.80. HRMS-ESI: *m*/*z* [M]^−^ calcd for [C_10_H_16_NO_3_]^−^: 198.1136; found: 198.1148.

### Cell culture, preparation of cell extracts, and immunoblot analysis

The BT474, MDA-MB-231, MDA-MB-468, MCF10A, HMEC, B16, MALME-3M, DU145, SH-SY5Y, K 562, and MOLT4 cells lines were purchased from American Type Culture Collection (ATCC) (Manassas, VA USA). The KMS-11, KMS-18, JIM-3, and 4937 cell lines were kindly provided by Dr. Olga Guryanova, University of Florida Health Cancer Center and the WM793 cell line was kindly provided by Dr. W. Douglas Cress, Moffitt Cancer Center. Derivation of the HCI-001/LVM2/LR10 cell line is previously described (Wang, et al. 2019b, Wang, et al. 2019c). MCF10A cell lines stably expressing MYC were generated as detailed in previous reports (Law, et al. 2002, Wang, et al. 2019b, Wang, et al. 2019c), as was the generation of the 231/E-Cad and 231/E-Cad-GFP cells lines (Law, et al. 2013). Cell lysates were prepared, as described previously (Law, et al. 2002). Immunoblot analysis was performed utilizing the following antibodies purchased from Cell Signaling Technology (Beverly, MA USA) {Ac-H3[K9], #9649; Ac-H3[K17], #4353; AcK (Acetyl lysine), #9441; p-Akt Substrate (RXXS/T), #9614; p-Akt[T308], #13038; CDCP1, #13794; p-CDCP1[Y707], #13111; p-CDCP1[Y734], #9050; pCDCP1[Y743], #13093; pCDCP1[Y806], #13024; Cyclin B, #4135; p-ERK, #9101; p-FAK[Y397], #8556; PD-L1, #13684; Rb, #9309; p-Rb[S780], #9307; p-Rb[S795], #9301; pRb[S807/811], #9308; p-S6[S235/236], #2211; p-p70 S6 Kinase[T389], #9205; p-Src[Y416], #6943; p-Src[Y527], #2105; pSTAT3[Y705], #9131; pSTAT3[Y727], #9134}, from Santa Cruz Biotechnology (Santa Cruz, CA USA) {β-Actin, sc-47778; Cyclin D1, sc-450; Myc, sc-40 and sc-42; p70 S6 Kinase, sc-230; Src, sc-18; STAT3, sc-7179}, from Upstate Biotechnology (Lake Placid, NY USA) {MPM2, 05-368}, and from BD Transduction Laboratories (San Jose, CA USA) {E-Cadherin, #610182; PAI-1, #612024}.

### Statistical Analysis

#### Statistics

Statistical analysis was performed using Student’s *t*-test. Error bars represent standard deviation of the mean unless otherwise indicated and all P values are two-tailed. *P* values are documented in either the figures or legends. Points from cell culture studies plotted on bar or line graphs are the average of six or more replicates and are representative of three or more independent experiments. Quantitation of immunoblot results was performed by scanning the blots, inverting the images with Photoshop (San Jose, CA), and quantitating band intensities using ImageJ (NIH). Bar graphs of band intensities are presented as the average and error bars correspond to the standard error of the mean.

## RESULTS

### Tranexamic acid suppresses cancer cell viability in vitro and tumor growth in vivo

TA and the other compounds used or mentioned in the work described here are listed in Fig. 1A. MTT cell viability assays showed that TA suppressed the viability of a variety of cancer cell lines including melanoma (B16, WM793, and MALME-3M), breast (MDA-MB-468 and BT474), prostate (DU145), and neuroblastoma (SH-SY5Y) lines (Fig. 1B). DU145 was one of the more TA-sensitive lines and exhibited statistically significant TA-mediated viability suppression at the lowest TA concentration tested 0.25% w/v compared with the vehicle control (Fig. 1C). Studies of TA effects on the viability of leukemia, lymphoma, and multiple myeloma lines using the XTT assay showed concentration-dependent reductions in cancer cell viability (Fig. 1D). The reduction of cell viability in MTT/XTT assays may be partially due to suppression of cell replication since TA inhibited DNA synthesis as measured in thymidine incorporation assays employing the MDA-MB-468 and BT474 cancer lines (Fig. S1A). The MYC oncogene is frequently overexpressed in human cancer (Rothberg 1987, Saksela 1990, Shiu, et al. 1993). However, MYC overexpression renders cancer cells vulnerable to some therapeutics, resulting in collateral vulnerabilities amenable to synthetic lethal targeting approaches (Hsieh and Dang 2016, Lee, et al. 2019). Therefore, we examined if MYC overexpression increased the sensitivity of the MCF10A human breast epithelial cell line to TA. MYC overexpression enhanced the TA suppression of cell viability across an array of TA concentrations suggesting that MYC overexpression may be useful as a marker to identify TA-sensitive tumors (Fig. 1E).

We employed a previously described metastatic, Lapatinib-resistant, xenograft breast model, 012/LVM2/LR10 (Wang, et al. 2019a, Wang, et al. 2019b), to examine if TA reduces the growth of breast tumors in experimental animals (Fig. 1F, upper panel). Tumor growth studies revealed that 375 mg/kg TA suppressed the growth of 012/LVM2/LR10 tumors without evidence of treatment toxicity (Fig. 1F, lower panel). At the 12-day treatment endpoint, tumors in the TA treatment group were approximately one-ninth the size of the tumors in the vehicle group (Fig. 1G).

### TA suppresses S6K1 and STAT3 phosphorylation

We next sought to examine the mechanisms through which TA impacts tumor growth. TA may exhibit anticancer activity through effects on Plasmin activity and via Plasmin-independent actions (Dunbar, et al. 2000, O’Grady, et al. 1981, Stonelake, et al. 1997, Suojanen, et al. 2009, Tanaka, et al. 1982). Previous work indicated that Plasmin cleaves the transmembrane protein CUB Domain-Containing Protein 1 (CDCP1) to produce a fragment that exhibits differential protein-protein interactions and signaling functions compared with the full-length protein, and suggested that cleaved CDCP1 (cCDCP1) may contribute to tumor growth and progression (Casar, et al. 2014, Law, et al. 2016, Wright, et al. 2016). However, the ability of TA to suppress CDCP1 cleavage has not been examined. TA reduced CDCP1 cleavage in MDA-MB-231 breast cancer cells (Fig. 2A). Transforming Growth Factor-β (TGFβ) or the glucocorticoid dexamethasone, which block CDCP1 cleavage by upregulating Plasminogen Activator Inhibitor-1 (PAI-1) as reported previously (Law, et al. 2013, Law, et al. 2016), TA did not increase PAI-1 levels (Fig. 2A). cCDCP1 binds E-cadherin and interferes with its cell-cell adhesive functions, while the full-length protein does not (Law, et al. 2016). Blocking CDCP1 cleavage via TA administration may be useful on restoring the anti-invasive functions of E-cadherin. Since full-length CDCP1 does not bind E-cadherin, we do not expect enforced E-cadherin expression alter TA inhibition of CDCP1 cleavage positively or negatively. To confirm this, we also examined if enforced expression of E-cadherin or an E-Cadherin-Green Fluorescent Protein (GFP) fusion protein influences TA blockade of CDCP1 cleavage (Fig. 2B). TA (0.5%) decreased CDCP1 cleavage in MDA-MB-231 cells irrespective of E-cadherin overexpression and did not significantly impact CDCP1 tyrosine phosphorylation on multiple sites previously implicated in regulating downstream signaling proteins including Src (He, et al. 2010, Liu, et al. 2011). Consistent with this, TA did not alter Src phosphorylation on activating (Y416) or inhibitory (Y527) sites (Fig. 2B). FAK and CDCP1 exhibit reciprocal tyrosine phosphorylation patterns in which FAK is heavily phosphorylated in cells attached to substrata, while CDCP1 is heavily phosphorylated in cells growing in suspension (Wortmann, et al. 2011). TA did not alter FAK phosphorylation on the Y397 site that reflects FAK activation (Thamilselvan, et al. 2007). Both the full length, 135 kDa, and cleaved, 85 kDa, forms of CDCP1 exhibit tyrosine phosphorylation at multiple sites.

**Fig. 2:**
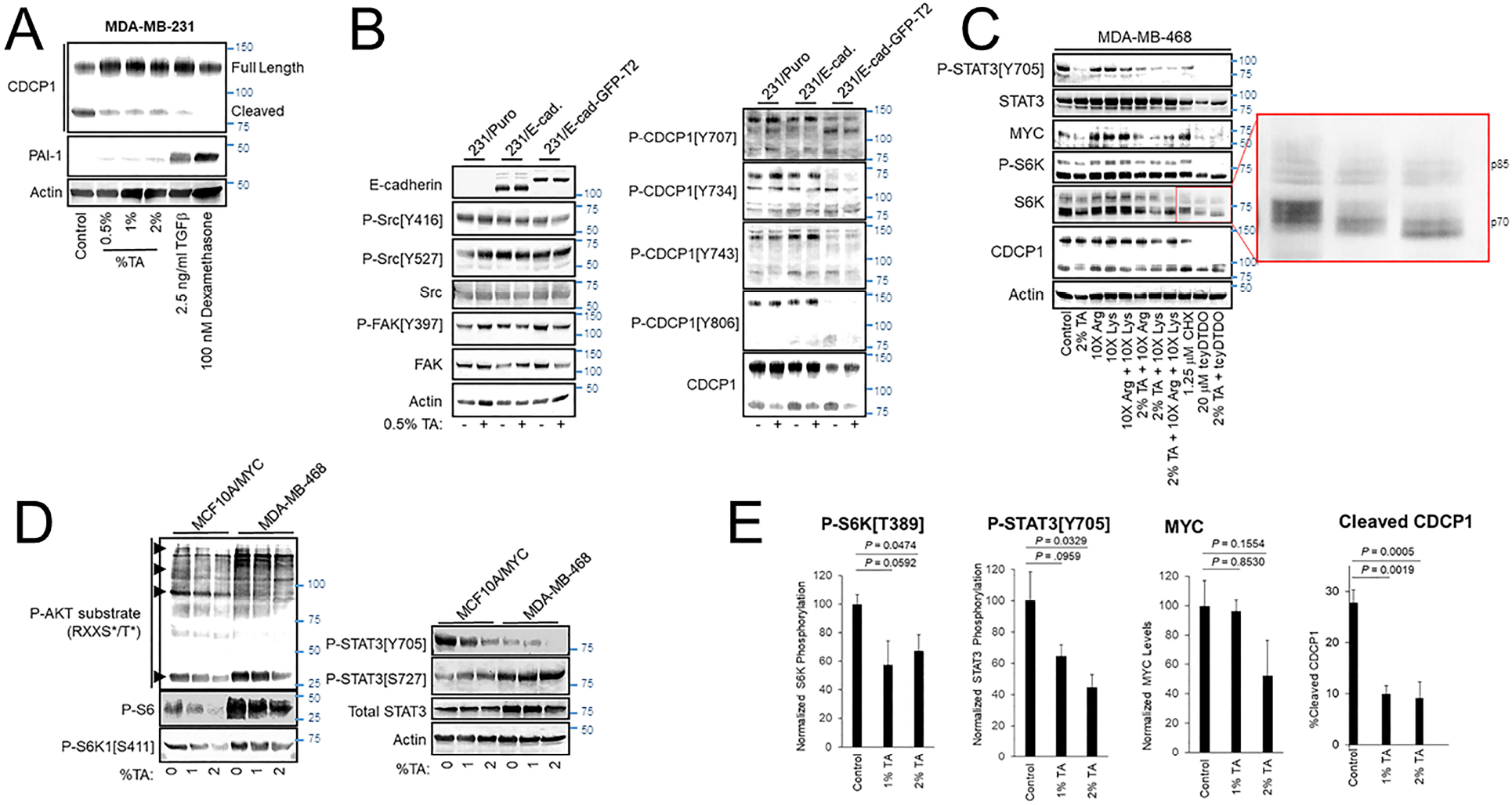
TA blocks CDCP1 cleavage, reduces protein synthesis and MYC expression, and decreases STAT3 and S6K1 phosphorylation. A. MDA-MB-231 cells were treated as indicated for 24 h and cell extracts were subjected to immunoblot analysis. B. Vector control MDA-MB-231 cells or cells overexpressing E-cadherin (E-Cad.) or an E-cadherin-Green Fluorescent Protein fusion (E-Cad-GFP) were treated for 24 h with vehicle or 0.5% TA and cell extracts were subjected to immunoblot analysis. C. MDA-MB-468 cells were treated for 24 h as indicated and subjected to immunoblot analysis. As indicated Lys and/or Arg were present in some treatments at 10X the concentration present in DMEM. The expanded red box shows altered mobility due to differential phosphorylation of the p70 and p85 isoforms of S6K1. Note that TA combined with tcyDTDO reduced S6K1 phosphorylation more than that observed with tcyDTDO alone. Inhibition of protein synthesis with 1.25 μM Cycloheximide (CHX) reduced STAT3 tyrosine phosphorylation, but did not alter MYC levels, S6K1 phosphorylation, or CDCP1 cleavage. D. Immunoblot analyses examining the effects of treating MYC overexpressing MCF10A cells or MDA-MB-468 cells for 24 h with 1 or 2% TA on STAT3 Ser and Tyr phosphorylation, S6K1 phosphorylation on Ser^411^, phosphorylation of the S6K1 substrate, S6, and phosphorylation of proteins on consensus phosphorylation sites for AKT. Note that the AKT consensus phosphorylation sequence overlaps with that of S6K1. E. Quantitation of the effects of 24 h treatment of MDA-MB-468 cells with 1% or 2% TA on S6K1 phosphorylation, STAT3 phosphorylation, MYC expression levels, or CDCP1 cleavage across multiple experiments.

We also screened the effects of TA on several other signaling pathways important in tumor growth and progression including the oncogenic transcription factors STAT3 and MYC, the mitogen-activated serine/threonine kinases S6 protein kinase (S6K), ERK, and Akt, and the levels and phosphorylation status of proteins that control cell cycle progression including Cyclin D1, Cyclin B, and Rb. TA reduced STAT3 tyrosine phosphorylation on its activating site, Y705, decreased MYC protein expression, and suppressed S6K1 phosphorylation on the activating site, T389 (Fig. 2C). TA had no effect on ERK, Akt, or Rb phosphorylation or Cyclin D1 or Cyclin B levels (Fig. S1B).

TA treatments were also carried out in the presence of 10X excesses of Lys or Arg to examine the ability of high levels of these amino acids to overcome TA effects. High Lys or Arg levels did not alter STAT3 phosphorylation, but increased S6K1 phosphorylation and MYC levels consistent with the sensitivity of S6K1 activation (Hidayat, et al. 2003, Iiboshi, et al. 1999) and MYC expression (Yue, et al. 2017) to amino acid levels (Fig. 2C). However, TA still reduced S6K1 phosphorylation and MYC levels in the presence of excess Lys or Arg. Treatment with the protein synthesis inhibitor cycloheximide (CHX) was included as a control to determine if inhibition of protein synthesis was sufficient to mimic any TA effects. CHX decreased STAT3 phosphorylation, but did not decrease MYC levels or S6K1 phosphorylation. Control protein synthesis experiments show that 1.25 μM CHX inhibits protein synthesis by approximately 60% (Fig. S1C).

Our group identified a class of compounds with anticancer activity termed disulfide bond disrupting agents (DDAs) that interfere with disulfide bond formation, induce endoplasmic reticulum (ER) stress, and inhibit protein synthesis (Ferreira, et al. 2015, Ferreira, et al. 2017, Wang, et al. 2019a, Wang, et al. 2019b). Results posted in a preprint suggest that DDAs act by inhibiting a subset of Protein Disulfide Isomerases (Law, et al. 2021). Therefore, we examined if DDAs induce overlapping responses with TA and therefore may be useful in combination regimens against cancer. The DDA tcyDTDO (Fig. 2C) decreased STAT3 and S6K1 phosphorylation and MYC levels.

Combined tcyDTDO/TA treatment reduced overall S6K1 phosphorylation more than either treatment alone as evidenced by electrophoretic mobility shifts of both the 70 and 85 kDa forms of S6K1 (Fig. 2C, expanded region). We next examined the effect of TA on the phosphorylation of STAT3 and S6K1 in the MCF10A cell line overexpressing MYC and the MDA-MB-468 breast cancer line. TA induced a concentration-dependent decrease in STAT3 phosphorylation on Y705 but had no effect on phosphorylation on S727 (Fig. 2D). The AGC-family kinases Akt and S6K1 have similar arginine-rich consensus substrate sequences. Immunoblot analyses using antibodies recognizing phospho-Akt substrates and the S6K1 substrate S6 showed a TA-induced decrease in phosphorylation at these sites. This was paralleled by reduced phosphorylation of the S6K1 site S411 that is also associated with kinase S6K1 activation (Han, et al. 1995). Together, the results presented thus far in Fig. 2 indicate that TA inhibits a subset of mitogenic/oncogenic signaling pathways. Quantification studies demonstrated that 2% w/v TA decreases phosphorylation of S6K1 on T389 and STAT3 on Y705, which are sites required for their kinase and transcriptional activities, respectively (Fig. 2E). TA reduced CDCP1 cleavage in a concentration-dependent and statistically significant manner (Fig. 2E). TA also decreased MYC expression levels, but this trend did not reach statistical significance.

### TA inhibits protein synthesis and reduces histone acetylation

Since TA and protein synthesis inhibition by CHX both decreased STAT3 tyrosine phosphorylation (Fig. 2C) and TA mimics Lys and Arg, we hypothesized that TA may inhibit protein synthesis. TA inhibited protein synthesis in a concentration-dependent manner, with 2% TA reducing protein synthesis by 60% (Fig. 3A). DDAs also inhibit protein synthesis (Ferreira, et al. 2017, Wang, et al. 2019b, Wang, et al. 2019c).Therefore, we examined if combining low concentrations of the DDA tcyDTDO and TA would suppress protein synthesis in an additive manner. Treatment with 1.25 μM tcyDTDO inhibited protein synthesis by 51% and addition of increasing concentrations of TA further reduced protein synthesis (Fig. 3B). Similarly, combining concentrations of TA and tcyDTDO that have little effect on cancer cell viability alone resulted in larger reductions in viability in MDA-MB-468 breast cancer cells (Fig. 3C).

**Fig. 3:**
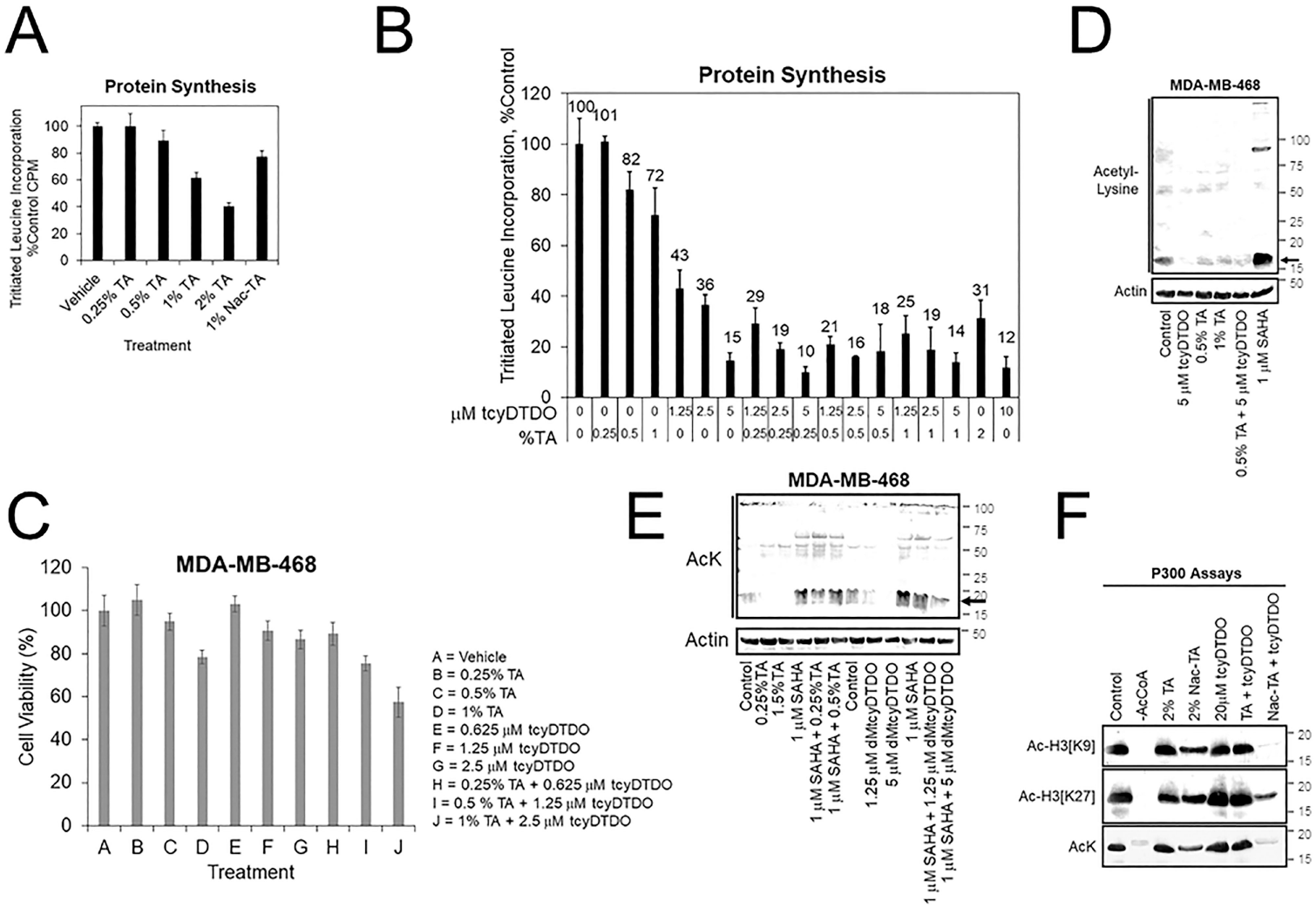
TA and DDAs exhibit partially overlapping anticancer mechanisms that include inhibition of protein synthesis, cell viability, and Histone acetylation. A. Protein synthesis assay, performed as in Fig. 2D, examining the effects of 24 h treatments with varying TA concentrations, or the *N*-acetylated TA metabolite NacTA, on protein synthesis in MDA-MB-468 cells. B. MDA-MB-468 cells were treated for 24 h with the indicated concentrations of tcyDTDO and TA and protein synthesis assays were performed. C. MTT cell viability assays performed after 72 h of the indicated treatments. D. MDA-MB-468 cells were treated for 24 h as indicated and cell extracts were analyzed by immunoblot with antibodies recognizing acetyl-lysine (AcK) and Actin as a loading control. SAHA, also referred to as Vorinostat, is a Histone Deacetylase (HDAC) inhibitor. E. MDA-MB-468 cells were treated for 24 h as indicated and cell extracts were analyzed by immunoblot with antibodies recognizing acetyl-lysine (AcK) and Actin as a loading control. The black arrow indicates the band corresponding to acetylated histones. F. *In vitro* acetyltransferase assays employing p300 as the acetyltransferase and Histone 3 as the substrate. Assays were performed for 60 min. at 37°C with the indicated additions and reaction mixtures were analyzed by immunoblot with the indicated antibodies recognizing acetyl-lysine or Histone 3 specifically acetylated on Lys residues 9 or 27. No acetylation was observed if Acetyl CoA was omitted from the reactions.

Suppression of histone acetylation by the p300/CBP enzymes is an emerging strategy for cancer therapy (Gu, et al. 2016, Lasko, et al. 2017, van Gils, et al. 2021, Wang, et al. 2017, Yang, et al. 2013). We hypothesized that TA may alter Lys acetylation through competitive inhibition of Lys acetyltransferases and next examined if TA affects acetylation of proteins on Lys. TA at 0.5 or 1% partially blocked protein Lys acetylation of a prominent ∼ 20 kDa band as detected with an acetyl-lysine antibody (Fig. 3D, arrow). This band is likely acetylated histones, which are the most abundant acetylated cellular proteins in this molecular size range (Hansen, et al. 2019). TcyDTDO also reduced acetylation of this band, and acetylation was strongly increased by treatment of the cells with the histone deacetylase inhibitor Vorinostat/SAHA. We next examined if TA could overcome the effects of SAHA. TA treatment reduced Lys acetylation, but TA did not override the increased acetylation induced by SAHA co-treatment (Fig. 3E). In contrast, treatment with the DDA dMtcyDTDO (Law, et al. 2021) decreased baseline Lys acetylation and blocked the SAHA-driven increase in acetylation.

The p300 enzyme is a major cellular histone acetyltransferase (HAT). We next performed HAT enzyme assays to determine if TA or tcyDTDO directly inhibit p300 activity. TA (2%) did not affect p300 acetylation of Histone 3 *in vitro* (Fig. 3F). TcyDTDO also did not directly inhibit p300 activity. TA is partly metabolized in the body by acetylation (0.5% of oral dose; http://www.accessdata.fda.gov/drugsatfda_docs/label/2020/022430s009lbl.pdf), and perhaps also in cultured cells. Therefore, we examined if the metabolite *N-*acetyl-TA (NacTA) alters p300 activity. NacTA partially inhibited p300 HAT activity and interestingly, combining NacTA and tcyDTDO strongly inhibited overall p300 activity (AcK), as well as acetylation of Histone H3 on Lys residues 9 and 27. Further studies are required to determine if TA, NacTA, or DDAs reduce cellular protein acetylation through effects on p300 or other HAT enzymes. However, the results in Fig. 3D-F suggest that TA and/or its metabolites and DDAs may mediate their effects in part by altering protein acetylation, which may alter gene expression through epigenetic mechanisms.

### Efficacy of TA/DDA mono- and combination anti-cancer therapy

The findings presented in Figs. 1-3 indicate that TA suppresses the growth of tumors derived from 012/LVM2/LR10 cells in a 12-day study, and cell culture studies show that TA may suppress cancer cell viability through multiple mechanisms. The results further suggest that TA mechanisms of anticancer action partially overlap with those of DDAs. Therefore, we examined the anticancer efficacy of 375 mg/kg TA, 10 mg/kg of the DDA dMtcyDTDO, or TA + dMtcyDTDO administered once daily by oral gavage in longer-term studies. Tumors were initiated by orthotopic injections of 10^6^ 012/LVM2/LR10 cells and treatment was initiated when tumors were detectable by palpation. By 20 days of treatment, tumors in vehicle-treated mice averaged over 200 mm^3^, while tumors in drug-treated animals were less than 50 mm^3^ (Fig. 4A). Immunoblot analysis of tumor extracts from each group showed that TA reduced S6K1 phosphorylation on Thr^389^, which is required for its kinase activity (Dennis, et al. 1998, McMahon, et al. 2002), to a greater extent on average than the other treatments (Fig. 4B). Quantification of the results in Fig. 4B indicated that TA suppressed S6K1 phosphorylation in a statistically significant manner (Fig. 4C). Rapamycin analogs (“rapalogs”) are in clinical use against human cancer and act in part by inhibiting S6K (Ferrari, et al. 1993, Meng and Zheng 2015), and S6K inhibitors are under development as cancer therapeutics (Nam, et al. 2019, Pearce, et al. 2010), consistent with a well-established role of S6K activation in the molecular pathogenesis of multiple cancers.

**Fig. 4:**
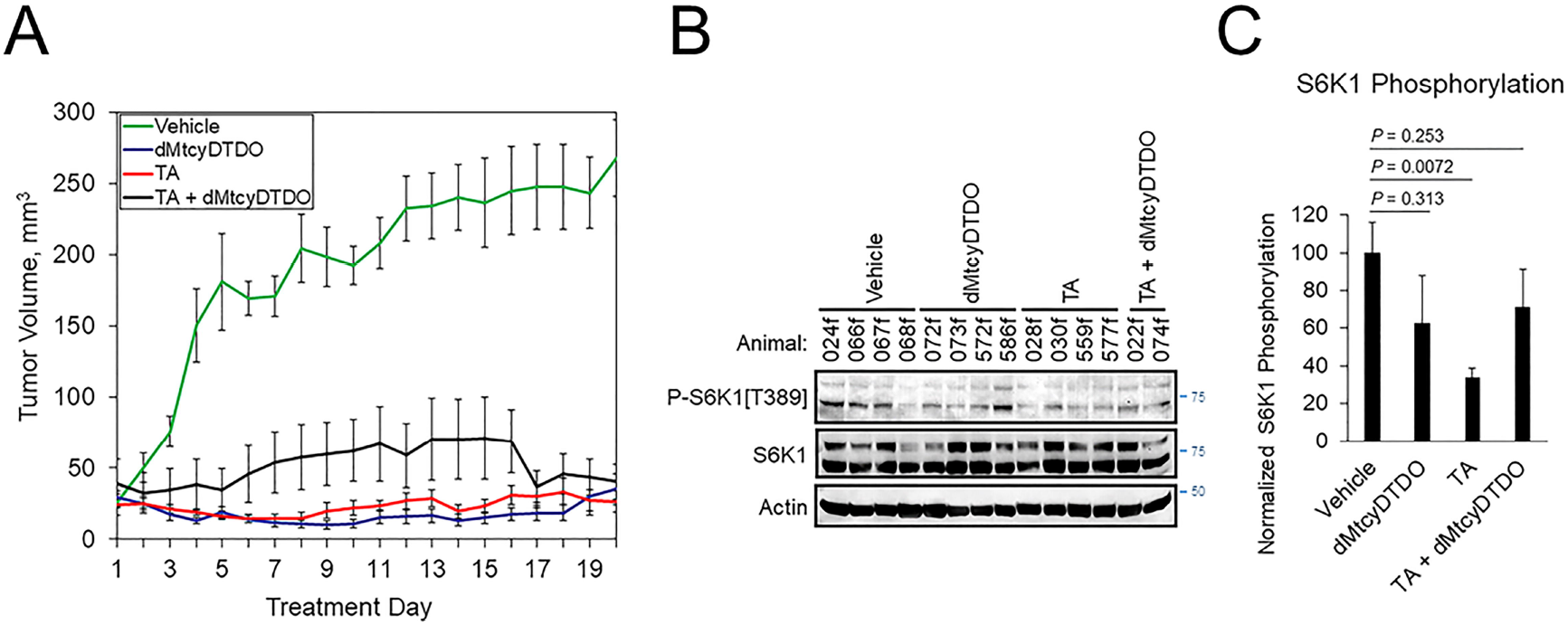
TA blockade of breast tumor growth is associated with reduced S6K1 phosphorylation. A. Growth curves of HCI-012/LVM2/LR10 tumors treated once daily by oral gavage with vehicle (peanut oil), 10 mg/kg dMtcyDTDO, 375 mg/kg TA or combined treatment with the 10 mg/kg dMtcyDTDO and 375 mg/kg TA. B. Immunoblot analysis of tumor extracts with the indicated antibodies. C. Plot of S6K1 phosphorylation normalized to S6K1 protein levels. Error bars represent Standard Error of the Mean. Pairwise statistical comparisons between the vehicle and drug treatment groups were made using Student’s *t*-test.

We previously observed that treatment with the DDA tcyDTDO caused widespread apoptosis of cancer cells in primary tumors and metastases without inducing apoptosis in adjacent normal tissues, resulting in substantial regions of dead tumor tissue (Wang, et al. 2019b). This indicates that estimating tumor volumes without taking into account viability of the cancer cells may underestimate the efficacy of anticancer regimens. Further, analysis of portions of such heterogeneous tumors may introduce sampling errors into downstream proteomic or transcriptomic analyses. Therefore, we examined the morphology of Hematoxylin and Eosin (H&E)-stained tumors from each of the four treatment groups. Two representative tumors from vehicle-treated mice (Fig. 5A) exhibited predominantly viable tumor tissue (T), with small regions of dead cancer cells (N). Higher magnification images showed mitotic figures, demonstrating cancer cell proliferation in the vehicle-treated tumors (yellow arrows). Representative tumors from dMtcyDTDO-treated animals in contrast, exhibited predominantly dead tumor tissue (N), with small areas of apparently viable tumor cells (T) (Fig. 5B). Closer inspection of the boundary between the dead and viable cancer cells showed tumor cells undergoing nuclear condensation and fragmentation (red arrows), consistent with apoptosis. Interestingly, tumors from TA-treated animals exhibited somewhat similar morphology to the tumors from dMtcyDTDO-treated animals with large regions of dead tumor tissue and tumor cells exhibiting nuclear condensation and fragmentation (red arrows, Fig. 5C). Tumors from mice treated with TA + dMtcyDTDO (Fig. 5D) exhibited similar morphology to the tumors from mice treated with TA or dMtcyDTDO with large regions of dead tumor tissue, tumor cells exhibiting nuclear condensation and fragmentation, and a lack of mitotic figures as observed in the tumors from vehicle-treated mice. A few previous studies have noted TA anticancer activity in xenograft models (Astedt and Trope 1980, Iwakawa and Tanaka 1982, Ogawa, et al. 1982), and in a case report (Astedt, et al. 1977), against several solid tumor types, including breast cancer. In these reports, TA anticancer activity was attributed to its antifibrinolytic actions. Our results demonstrate TA efficacy against breast tumors in animal models and identify several novel TA-induced biochemical responses that may contribute to TA anticancer activity.

**Fig. 5:**
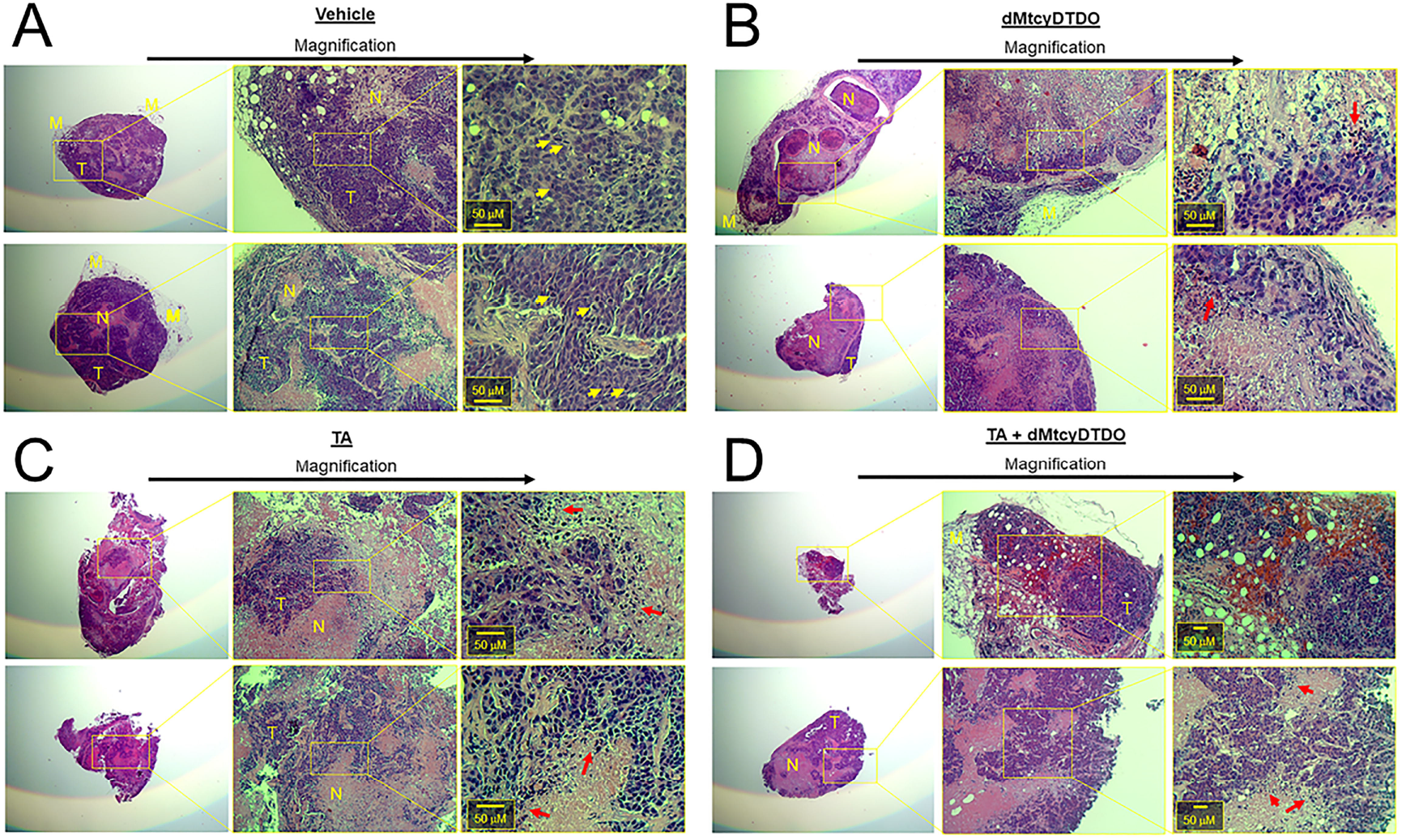
TA and dMtcyDTDO cause tumor cell death in animal models. At endpoint, two tumors from the A. Vehicle, B. dMtcyDTDO, C. TA, and D. TA + dMtcyDTDO treatment groups were paraffin-embedded, sectioned and stained with hematoxylin and eosin (H&E). Each section is shown at three different magnifications with scale bars on the highest magnification images. T refers to viable tumor tissue, N, necrotic/dead tissue, and M, mammary tissue. Yellow arrows denote cells undergoing mitosis and red arrows indicate cells undergoing nuclear condensation and fragmentation, indicative of cell death.

## DISCUSSION

Previous work suggested that TA might exhibit anticancer activity by inhibiting Plasmin activity and by preventing Plasminogen conversion to Plasmin (Dunbar, et al. 2000, O’Grady, et al. 1981, Stonelake, et al. 1997, Suojanen, et al. 2009, Tanaka, et al. 1982). The results presented here indicate that TA blocks cleavage of the Plasmin substrate CDCP1. Some reports indicate that the cleaved form of CDCP1 exhibits enhanced oncogenic functions and a different spectrum of protein-protein interactions than the full-length protein (Casar, et al. 2014, Law, et al. 2016, Wright, et al. 2016). Suppression of CDCP1 cleavage by TA may be preferable to that induced by TGFβ and glucocorticoids since these agents block Plasminogen activation by upregulating Plasminogen Activator Inhibitor-1 (PAI-1) and activate transcriptional programs that may not be beneficial in the cancer setting. However, blockade of Plasminogen activation and cleavage of CDCP1 and other Plasmin substrates represents only one of the biochemical actions of TA that could contribute to its anticancer activity.

The MYC and STAT3 oncoproteins are widely activated across human cancers and the mTORC1/S6K1 pathway is frequently stimulated in malignancies. Of interest is the observation that MYC transformed MCF10A mammary epithelial cells are more sensitive to TA-mediated reduction in cell viability than vector control cells. This may be due to MYC-mediated monopolization of the protein translation machinery and regulation of the translation factors eIF4E, eIF4A, and eIF4G (Flynn and Hogarty 2018, Ruggero 2009) and could explain in part how TA suppresses breast tumor growth in animal models without evidence of toxicity. Given these observations, more work is warranted to elucidate in detail the biochemical mechanisms by which TA suppresses MYC expression, and STAT3 and S6K1 phosphorylation, and reduces histone acetylation. The protein synthesis inhibitor Cycloheximide partially blocked STAT3 tyrosine phosphorylation (Fig. 2C), but did not reduce MYC levels or S6K1 phosphorylation, indicating that the latter two TA effects are independent of protein synthesis inhibition.

TA inhibition of protein synthesis is of particular interest because Asparaginase, which depletes asparagine and glutamine, is in clinical use against leukemia (reviewed in (Nikolaev 1969)) and functions in part by suppressing protein synthesis (Benvenisty 1972, Ellem, et al. 1970, Kessel and Bosmann 1970) and inhibiting mTORC1/S6K1 signaling (Nikonorova, et al. 2018).Asparaginase resistance remains a significant clinical problem (Apfel, et al. 2021) and as a protein drug, Asparaginase suffers from issues associated with instability and rapid clearance (Varshosaz and Anvari 2018) and allergenicity (Belen, et al. 2019). Thus, a small molecule drug such as TA or a TA analog may be useful against Asparaginase-resistant malignancies or in patients with Asparaginase allergies. Due to the similarity of TA to Lys/Arg, future studies should examine the possibility that TA suppresses protein synthesis by competing with Lys/Arg for plasma membrane amino acid transporters or by inhibiting charging of Lys or Arg tRNAs in the cytoplasm. Either of these actions could explain the TA-induced suppression of protein synthesis and S6K1 phosphorylation (Drummond, et al. 2008).

Results posted in a preprint suggest that DDAs may suppress tumor growth by inhibiting a subset of Protein Disulfide Isomerases (PDIs) (Law, et al. 2021). Interestingly, DDAs exhibit some activities that overlap with those of TA, including decreased MYC expression, sensitivity enhancement by MYC overexpression, lowering of STAT3 tyrosine phosphorylation and S6K1 phosphorylation on Thr^389^ and Ser^411^, inhibition of protein synthesis, and reduced of histone acetylation. Fig. S1D summarizes the common biochemical responses to TA and DDA that are hypothesized to mediate in part, the anticancer activity of these compounds. The results in Fig. 4 examining TA/DDA mono- and combination regimens against breast tumors indicate that TA suppresses S6K1 phosphorylation on Thr^389^ in tumors *in vivo*.

TA exhibits anticancer activity in mice at a dosage of 375 mg/kg (Figs. 1F, 4A). Future studies are required to determine whether similar TA anticancer activity is observed at lower dosages. Clinically, TA has been used at dosages of 20 mg/kg to suppress postoperative bleeding (Andersson, et al. 1968). The most common intravenous dosages used clinically are 10 to 20 mg/kg, given every six to eight hours, or simply before and after surgery (Lecker, et al. 2012). Administration of 240 mg/h TA was used to prevent life-threatening bleeding in advanced cancer patients (Atreya 2021). However, dosages as high as 100 mg/kg followed by 10 mg/kg/hour were given previously, until it was found that total doses of 80-100 mg led to a small increase in post-operative seizures. An important toxicity of TA when used at high dosages is seizures (Kalavrouziotis, et al. 2012, Mohseni, et al. 2009). This is thought to result primarily from TA induction of neuronal hyperexcitability due to antagonism of glycine binding to glycine receptors, which hyperpolarize neurons (Lecker, et al. 2012). Importantly, this neuronal hyperexcitability may be overcome using the common surgical anesthetics isoflurane and propofol (Lecker, et al. 2012). TA side effects may also be avoided by using it against skin cancers via topical administration or against other surface-accessible malignancies such as superficial bladder cancers (Javadpour 1989) or lung tumors (Jang, et al. 2006, Moghissi, et al. 2004).

It should be noted that while TA is not currently approved for use as an anticancer agent, TA is employed with increasing frequency in various aspects of the clinical cancer management. TA is often used to limit bleeding during surgical tumor resections (Jaffer, et al. 2021, Longo, et al. 2018, Wu, et al. 2006). TA treatment is associated with a number of malignant cells and the generation of ascites fluid in patients with disseminated ovarian cancer (Kikuchi, et al. 1986, Soma, et al. 1980). In the treatment of node-positive, operative breast cancer, the axillary lymph nodes are dissected out, and this is associated with excessive axillary drainage and risk of infection. TA treatment reduces axillary drainage and decreases the rate of complications in these patients (Lohani, et al. 2020, Lohani, et al. 2021, Oertli, et al. 1994). It will be important to determine in these clinical scenarios where TA is used to control treatment-associated morbidities, if TA also contributes to improved patient outcome through direct actions against any remaining cancer cells. If so, then in cancers such as Triple-Negative Breast Cancer (TNBC), which exhibit high rates of post-surgical recurrence (Pilewskie, et al. 2014, Radosa, et al. 2017), TA treatment before or after tumor resection might reduce the rate of cancer recurrence.

The potential for beneficial actions of TA in inhibiting Plasmin activation and activity in the cancer setting are well appreciated, but to our knowledge, the present study is the first to survey the effects of TA on an array of other signaling pathways and proteins with roles in tumor growth and progression. Given the established ability of TA to antagonize Lys and Gly binding sites, and the likelihood that TA also antagonizes arginine and histidine binding sites due to its positively charged amine group may explain the different pharmacological mechanisms of action exhibited by TA. Additional work is needed to elucidate how TA mediates its Plasmin-independent signaling effects. Additionally, TA is anti-inflammatory and modulates the innate immune system (Draxler, et al. 2019). This study by Draxler et al was performed alongside the major prospective safety study of TA, ATACAS, where they found that use of TA did not cause an increase in thromboembolic events in the only prospective study with sufficient power to prove this point. The Draxler adjunctive study found that the patients who received TA had significantly fewer post-operative infections, despite the fact that every patient received prophylactic antibiotics.

We hypothesize that the multiple anticancer activities of TA may delay or block the acquisition of tumor resistance, and this might be further enhanced by combining TA with other classes of anticancer agents and/or therapies. It will also be important in future studies to determine if TA metabolites such as NacTA play significant roles in the responses of cancer cells to TA. The observation that TA elicits multiple biochemical responses that are expected to block tumor growth and progression suggests that it may be possible to design new TA analogs that more potently or selectively trigger individual downstream TA anticancer actions.

## ACKNOWLEDGEMENTS

This study was funded in part by NIH/NCI grant R21CA252400 and Tranexamic Technologies, LLC (BL and RC). Work performed by CDH was supported in part by NIH/NINDS grant R01NS102624. OG was supported by R01DK121831. We thank Drs. Daiqing Liao (UF) and Matthew Waddell (UF) for advice on p300 acetyltransferase assays and Dr. Michael Kilberg (UF) for providing the MOLT4s cell line. We are grateful to Dr. W. Douglas Cress (Moffitt Cancer Center) for providing the WM793 cell line.

## Author Contributions

Conceptualized Studies: ML, AG, MW, RC, BL

Performed Experiments: ML, BD, AG, MW, BL

Synthesized Compounds: EY, AG, ZD

Analyzed Results: ML, BD, MW, CDH, BL

Provided Critical Reagents: AG, OG, CDH, RC

Participated in Manuscript Writing/Editing: ML, BD, AG, MW, OG, CDH, RC, BL

Obtained Funding for the Study: BL, RC, CDH

## REFERENCES

Andersson L, Nilsoon IM, Colleen S, Granstrand B, Melander B. 1968. Role of urokinase and tissue activator in sustaining bleeding and the management thereof with EACA and AMCA. Ann N Y Acad Sci. Jun 28;146:642–658.

Apfel V, Begue D, Cordo V, Holzer L, Martinuzzi L, Buhles A, et al. 2021. Therapeutic Assessment of Targeting ASNS Combined with l-Asparaginase Treatment in Solid Tumors and Investigation of Resistance Mechanisms. ACS Pharmacol Transl Sci. Feb 12;4:327–337.

Astedt B, Mattsson W, Trope C. 1977. Treatment of advanced breast cancer with chemotherapeutics and inhibition of coagulation and fibrinolysis. Acta Med Scand.201:491–493.

Astedt B, Trope C. 1980. Effect of tranexamic acid on progress of experimental tumours and on DNA-synthesis. Experientia. Jun 15;36:679–680.

Atreya S. 2021. High-dose Continuous Infusion of Tranexamic Acid for Controlling Life-threatening Bleed in Advanced Cancer Patients. Indian J Palliat Care. Jan-Mar;27:172–175.

Belen LH, Lissabet JB, de Oliveira Rangel-Yagui C, Effer B, Monteiro G, Pessoa A, et al. 2019. A structural in silico analysis of the immunogenicity of l-asparaginase from Escherichia coli and Erwinia carotovora. Biologicals. May;59:47–55.

Benvenisty DS. 1972. [Inhibition of synthesis of protein and nucleic acid in mouse and human leukemia cells by L-asparaginase in vitro]. Harefuah. Jan 16;82:68–70.

Besse L, Besse A, Mendez-Lopez M, Vasickova K, Sedlackova M, Vanhara P, et al. 2019. A metabolic switch in proteasome inhibitor-resistant multiple myeloma ensures higher mitochondrial metabolism, protein folding and sphingomyelin synthesis. Haematologica. Sep;104:e415–e419.

Blee TK, Gray NK, Brook M. 2015. Modulation of the cytoplasmic functions of mammalian post-transcriptional regulatory proteins by methylation and acetylation: a key layer of regulation waiting to be uncovered? Biochem Soc Trans. Dec;43:1285–1295.

Casar B, Rimann I, Kato H, Shattil SJ, Quigley JP, Deryugina EI. 2014. In vivo cleaved CDCP1 promotes early tumor dissemination via complexing with activated beta1 integrin and induction of FAK/PI3K/Akt motility signaling. Oncogene. Jan 9;33:255-268. Epub 2012/12/05.

Corsino P, Horenstein N, Ostrov D, Rowe T, Law M, Barrett A, et al. 2009. A novel class of cyclin-dependent kinase inhibitors identified by molecular docking act through a unique mechanism. J Biol Chem. Oct 23;284:29945-29955. Epub 2009/08/28.

Dennis PB, Pullen N, Pearson RB, Kozma SC, Thomas G. 1998. Phosphorylation sites in the autoinhibitory domain participate in p70(s6k) activation loop phosphorylation. J Biol Chem. Jun 12;273:14845–14852.

Draxler DF, Awad MM, Hanafi G, Daglas M, Ho H, Keragala CB, et al. 2019. Tranexamic acid influences the immune response, but not bacterial clearance in a model of post-traumatic brain injury pneumonia. J Neurotrauma. May 29.

Drummond MJ, Bell JA, Fujita S, Dreyer HC, Glynn EL, Volpi E, et al. 2008. Amino acids are necessary for the insulin-induced activation of mTOR/S6K1 signaling and protein synthesis in healthy and insulin resistant human skeletal muscle. Clin Nutr. Jun;27:447–456.

Dunbar SD, Ornstein DL, Zacharski LR. 2000. Cancer treatment with inhibitors of urokinase-type plasminogen activator and plasmin. Expert Opin Investig Drugs. Sep;9:2085–2092.

Ellem KA, Fabrizio AM, Jackson L. 1970. The dependence of DNA and RNA synthesis on protein synthesis in asparaginase-treated lymphoma cells. Cancer Res. Feb;30:515–527.

Ferrari S, Pearson RB, Siegmann M, Kozma SC, Thomas G. 1993. The immunosuppressant rapamycin induces inactivation of p70s6k through dephosphorylation of a novel set of sites. J Biol Chem. Aug 5;268:16091–16094.

Ferreira RB, Law ME, Jahn SC, Davis BJ, Heldermon CD, Reinhard M, et al. 2015. Novel agents that downregulate EGFR, HER2, and HER3 in parallel. Oncotarget. Apr 30;6:10445–10459. Epub 2015/04/14.

Ferreira RB, Wang M, Law ME, Davis BJ, Bartley AN, Higgins PJ, et al. 2017. Disulfide bond disrupting agents activate the unfolded protein response in EGFR- and HER2-positive breast tumor cells. Oncotarget. Apr 25;8:28971–28989.

Flynn AT, Hogarty MD. 2018. Myc, Oncogenic Protein Translation, and the Role of Polyamines. Med Sci (Basel). May 25;6.

Gu ML, Wang YM, Zhou XX, Yao HP, Zheng S, Xiang Z, et al. 2016. An inhibitor of the acetyltransferases CBP/p300 exerts antineoplastic effects on gastrointestinal stromal tumor cells. Oncol Rep. Nov;36:2763–2770.

Han JW, Pearson RB, Dennis PB, Thomas G. 1995. Rapamycin, wortmannin, and the methylxanthine SQ20006 inactivate p70s6k by inducing dephosphorylation of the same subset of sites. J Biol Chem. Sep 8;270:21396–21403.

Hansen BK, Gupta R, Baldus L, Lyon D, Narita T, Lammers M, et al. 2019. Analysis of human acetylation stoichiometry defines mechanistic constraints on protein regulation. Nat Commun. Mar 5;10:1055.

He Y, Wortmann A, Burke LJ, Reid JC, Adams MN, Abdul-Jabbar I, et al. 2010. Proteolysis-induced N-terminal ectodomain shedding of the integral membrane glycoprotein CUB domain-containing protein 1 (CDCP1) is accompanied by tyrosine phosphorylation of its C-terminal domain and recruitment of Src and PKCdelta. J Biol Chem. Aug 20;285:26162-26173. Epub 2010/06/17.

Hidayat S, Yoshino K, Tokunaga C, Hara K, Matsuo M, Yonezawa K. 2003. Inhibition of amino acid-mTOR signaling by a leucine derivative induces G1 arrest in Jurkat cells. Biochem Biophys Res Commun. Feb 7;301:417–423.

Hsieh AL, Dang CV. 2016. MYC, Metabolic Synthetic Lethality, and Cancer. Recent Results Cancer Res.207:73–91.

Iiboshi Y, Papst PJ, Kawasome H, Hosoi H, Abraham RT, Houghton PJ, et al. 1999. Amino acid-dependent control of p70(s6k). Involvement of tRNA aminoacylation in the regulation. J Biol Chem. Jan 8;274:1092–1099.

Iwakawa A, Tanaka K. 1982. Effect of fibrinolysis inhibitor and chemotherapeutics on the growth of human cancers transplanted into nude mice and in tissue culture. Invasion Metastasis.2:232–248.

Iwamoto M. 1975. Plasminogen-plasmin system IX. Specific binding of tranexamic acid to plasmin. Thromb Diath Haemorrh. Jun 30;33:573–585.

Izzo A, Schneider R. 2010. Chatting histone modifications in mammals. Brief Funct Genomics. Dec;9:429–443.

Jaffer AA, Karanicolas PJ, Davis LE, Behman R, Hanna SS, Law CH, et al. 2021. The impact of tranexamic acid on administration of red blood cell transfusions for resection of colorectal liver metastases. HPB (Oxford). Feb;23:245–252.

Jang TW, Kim HK, Oak CH, Jung MH. 2006. Photodynamic therapy in early lung cancer: a report of two cases. Korean J Intern Med. Sep;21:178–182.

Javadpour N. 1989. Photodynamic therapy of superficial bladder cancer. Md Med J. Jun;38:461.

Kalavrouziotis D, Voisine P, Mohammadi S, Dionne S, Dagenais F. 2012. High-dose tranexamic acid is an independent predictor of early seizure after cardiopulmonary bypass. Ann Thorac Surg. Jan;93:148–154.

Kessel D, Bosmann HB. 1970. Effects of L-asparaginase on protein and glycoprotein synthesis. FEBS Lett. Sep 24;10:85–88.

Kikuchi Y, Kizawa I, Oomori K, Matsuda M, Kato K. 1986. Adjuvant effects of tranexamic acid to chemotherapy in ovarian cancer patients with large amount of ascites. Acta Obstet Gynecol Scand.65:453–456.

Kusakabe T, Yamazaki, K., Ohgiya, T., and Shibuya, K. 2010. Method for preparing trans-{4-[(alkylamino) methyl]-cyclohexyl}acetic acid ester. https://patentsgooglecom/patent/US8841478B2/en.

Laskey RA, Gorlich D, Madine MA, Makkerh JP, Romanowski P. 1996. Regulatory roles of the nuclear envelope. Exp Cell Res. Dec 15;229:204–211.

Lasko LM, Jakob CG, Edalji RP, Qiu W, Montgomery D, Digiammarino EL, et al. 2017. Discovery of a selective catalytic p300/CBP inhibitor that targets lineage-specific tumours. Nature. Oct 5;550:128–132.

Law BK, Chytil A, Dumont N, Hamilton EG, Waltner-Law ME, Aakre ME, et al. 2002. Rapamycin potentiates transforming growth factor beta-induced growth arrest in nontransformed, oncogene-transformed, and human cancer cells. Mol Cell Biol. Dec;22:8184-8198. Epub 2002/11/06.

Law BK, Norgaard P, Moses HL. 2000a. Farnesyltransferase inhibitor induces rapid growth arrest and blocks p70s6k activation by multiple stimuli. J Biol Chem. Apr 14;275:10796-10801. Epub 2001/02/07.

Law BK, Waltner-Law ME, Entingh AJ, Chytil A, Aakre ME, Norgaard P, et al. 2000b. Salicylate-induced growth arrest is associated with inhibition of p70s6k and down-regulation of c-myc, cyclin D1, cyclin A, and proliferating cell nuclear antigen. J Biol Chem. Dec 8;275:38261-38267. Epub 2000/09/20.

Law ME, Corsino PE, Jahn SC, Davis BJ, Chen S, Patel B, et al. 2013. Glucocorticoids and histone deacetylase inhibitors cooperate to block the invasiveness of basal-like breast cancer cells through novel mechanisms. Oncogene. Mar 7;32:1316-1329. Epub 2012/05/01.

Law ME, Ferreira RB, Davis BJ, Higgins PJ, Kim JS, Castellano RK, et al. 2016. CUB domain-containing protein 1 and the epidermal growth factor receptor cooperate to induce cell detachment. Breast Cancer Res.18:80.

Law ME, Yaaghubi E, Ghilardi AF, Davis BJ, Ferreira RB, Koh J, et al. 2021. Inhibitors of ERp44, PDIA1, and AGR2 induce disulfide-mediated oligomerization of Death Receptors 4 and 5 and cancer cell death. bioRxiv.2021.2001.2013.426390.

Lecker I, Wang DS, Romaschin AD, Peterson M, Mazer CD, Orser BA. 2012. Tranexamic acid concentrations associated with human seizures inhibit glycine receptors. J Clin Invest. Dec;122:4654–4666.

Lee HY, Cha J, Kim SK, Park JH, Song KH, Kim P, et al. 2019. c-MYC Drives Breast Cancer Metastasis to the Brain, but Promotes Synthetic Lethality with TRAIL. Mol Cancer Res. Feb;17:544–554.

Liu H, Ong SE, Badu-Nkansah K, Schindler J, White FM, Hynes RO. 2011. CUB-domain-containing protein 1 (CDCP1) activates Src to promote melanoma metastasis. Proc Natl Acad Sci U S A. Jan 25;108:1379-1384. Epub 2011/01/12.

Lohani KR, Kumar C, Kataria K, Srivastava A, Ranjan P, Dhar A. 2020. Role of tranexamic acid in axillary lymph node dissection in breast cancer patients. Breast J. Jul;26:1316–1320.

Lohani KR, Kumar C, Kataria K, Srivastava A, Ranjan P, Dhar A. 2021. Role of tranexamic acid in axillary lymph node dissection in breast cancer patients: Does it help in reducing lymphedema? Breast J. May;27:502.

Longo MA, Cavalheiro BT, de Oliveira Filho GR. 2018. Systematic review and meta-analyses of tranexamic acid use for bleeding reduction in prostate surgery. J Clin Anesth. Aug;48:32–38.

McMahon LP, Choi KM, Lin TA, Abraham RT, Lawrence JC, Jr. 2002. The rapamycin-binding domain governs substrate selectivity by the mammalian target of rapamycin. Mol Cell Biol. Nov;22:7428–7438.

Meng LH, Zheng XF. 2015. Toward rapamycin analog (rapalog)-based precision cancer therapy. Acta Pharmacol Sin. Oct;36:1163–1169.

Moghissi K, Dixon K, Thorpe JA, Oxtoby C, Stringer MR. 2004. Photodynamic therapy (PDT) for lung cancer: the Yorkshire Laser Centre experience. Photodiagnosis Photodyn Ther. Nov;1:253–262.

Mohseni K, Jafari A, Nobahar MR, Arami A. 2009. Polymyoclonus seizure resulting from accidental injection of tranexamic acid in spinal anesthesia. Anesth Analg. Jun;108:1984–1986.

Nam KH, Yi SA, Nam G, Noh JS, Park JW, Lee MG, et al. 2019. Identification of a novel S6K1 inhibitor, rosmarinic acid methyl ester, for treating cisplatin-resistant cervical cancer. BMC Cancer. Aug 6;19:773.

Nikolaev A. 1969. [The use of asparaginase preparations for the treatment of tumors and leukemia (review of the literature)]. Probl Gematol Pereliv Krovi. Jul;14:51–60.

Nikonorova IA, Mirek ET, Signore CC, Goudie MP, Wek RC, Anthony TG. 2018. Time-resolved analysis of amino acid stress identifies eIF2 phosphorylation as necessary to inhibit mTORC1 activity in liver. J Biol Chem. Apr 6;293:5005–5015.

O’Grady RL, Upfold LI, Stephens RW. 1981. Rat mammary carcinoma cells secrete active collagenase and activate latent enzyme in the stroma via plasminogen activator. Int J Cancer. Oct 15;28:509–515.

Oertli D, Laffer U, Haberthuer F, Kreuter U, Harder F. 1994. Perioperative and postoperative tranexamic acid reduces the local wound complication rate after surgery for breast cancer. Br J Surg. Jun;81:856–859.

Ogawa H, Sekiguchi F, Tanaka N, Ono K, Tanaka K, Kinjo M, et al. 1982. Effect of antifibrinolysis treatment on human cancer in nude mice. Anticancer Res. Nov-Dec;2:339–344.

Pearce LR, Alton GR, Richter DT, Kath JC, Lingardo L, Chapman J, et al. 2010. Characterization of PF-4708671, a novel and highly specific inhibitor of p70 ribosomal S6 kinase (S6K1). Biochem J. Oct 15;431:245–255.

Pilewskie M, Ho A, Orell E, Stempel M, Chen Y, Eaton A, et al. 2014. Effect of margin width on local recurrence in triple-negative breast cancer patients treated with breast-conserving therapy. Ann Surg Oncol. Apr;21:1209–1214.

Radosa JC, Eaton A, Stempel M, Khander A, Liedtke C, Solomayer EF, et al. 2017. Evaluation of Local and Distant Recurrence Patterns in Patients with Triple-Negative Breast Cancer According to Age. Ann Surg Oncol. Mar;24:698–704.

Rothberg PG. 1987. The role of the oncogene c-myc in sporadic large bowel cancer and familial polyposis coli. Semin Surg Oncol.3:152–158.

Ruggero D. 2009. The role of Myc-induced protein synthesis in cancer. Cancer Res. Dec 1;69:8839–8843.

Saksela K. 1990. myc genes and their deregulation in lung cancer. J Cell Biochem. Mar;42:153–180.

Shiu RP, Watson PH, Dubik D. 1993. c-myc oncogene expression in estrogen-dependent and -independent breast cancer. Clin Chem. Feb;39:353–355.

Soma H, Sashida T, Yoshida M, Miyashita T, Nakamura A. 1980. Treatment of advanced ovarian cancer with fibrinolytic inhibitor (tranexamic acid). Acta Obstet Gynecol Scand.59:285–287.

Stonelake PS, Jones CE, Neoptolemos JP, Baker PR. 1997. Proteinase inhibitors reduce basement membrane degradation by human breast cancer cell lines. Br J Cancer.75:951–959.

Suojanen J, Sorsa T, Salo T. 2009. Tranexamic acid can inhibit tongue squamous cell carcinoma invasion in vitro. Oral Dis. Mar;15:170–175.

Tanaka N, Ogawa H, Kinjo M, Kohga S, Tanaka K. 1982. Ultrastructural study of the effects of tranexamic acid and urokinase on metastasis of Lewis lung carcinoma. Br J Cancer. Sep;46:428–435.

Thamilselvan V, Craig DH, Basson MD. 2007. FAK association with multiple signal proteins mediates pressure-induced colon cancer cell adhesion via a Src-dependent PI3K/Akt pathway. FASEB J. Jun;21:1730–1741.

Turner BM. 2000. Histone acetylation and an epigenetic code. Bioessays. Sep;22:836–845.

Tweedie-Cullen RY, Brunner AM, Grossmann J, Mohanna S, Sichau D, Nanni P, et al. 2012. Identification of combinatorial patterns of post-translational modifications on individual histones in the mouse brain. PLoS One.7:e36980.

van Gils N, Martianez Canales T, Vermue E, Rutten A, Denkers F, van der Deure T, et al. 2021. The Novel Oral BET-CBP/p300 Dual Inhibitor NEO2734 Is Highly Effective in Eradicating Acute Myeloid Leukemia Blasts and Stem/Progenitor Cells. Hemasphere. Aug;5:e610.

Varshosaz J, Anvari N. 2018. Enhanced stability of L-asparaginase by its bioconjugation to poly(styrene-co-maleic acid) and Ecoflex nanoparticles. IET Nanobiotechnol. Jun;12:466–472.

Wang M, Ferreira RB, Law ME, Davis BJ, Yaaghubi E, Ghilardi AF, et al. 2019a. A novel proteotoxic combination therapy for EGFR+ and HER2+ cancers. Oncogene. May;38:4264–4282.

Wang M, Law ME, Davis BJ, Yaaghubi E, Ghilardi AF, Ferreira RB, et al. 2019b. Disulfide bond-disrupting agents activate the tumor necrosis family-related apoptosis-inducing ligand/death receptor 5 pathway. Cell Death Discov.5:153.

Wang MX, Ferreira RB, Law ME, Davis BJ, Yaaghubi E, Ghilardi AF, et al. 2019c. A novel proteotoxic combination therapy for EGFR+ and HER2+cancers. Oncogene. May 30;38:4264–4282.

Wang YM, Gu ML, Meng FS, Jiao WR, Zhou XX, Yao HP, et al. 2017. Histone acetyltransferase p300/CBP inhibitor C646 blocks the survival and invasion pathways of gastric cancer cell lines. Int J Oncol. Dec;51:1860–1868.

Wortmann A, He Y, Christensen ME, Linn M, Lumley JW, Pollock PM, et al. 2011. Cellular settings mediating Src Substrate switching between focal adhesion kinase tyrosine 861 and CUB-domain-containing protein 1 (CDCP1) tyrosine 734. J Biol Chem. Dec 9;286:42303-42315. Epub 2011/10/14.

Wright HJ, Arulmoli J, Motazedi M, Nelson LJ, Heinemann FS, Flanagan LA, et al. 2016. CDCP1 cleavage is necessary for homodimerization-induced migration of triple-negative breast cancer. Oncogene. Sep 08;35:4762–4772.

Wu CC, Ho WM, Cheng SB, Yeh DC, Wen MC, Liu TJ, et al. 2006. Perioperative parenteral tranexamic acid in liver tumor resection: a prospective randomized trial toward a “blood transfusion”-free hepatectomy. Ann Surg. Feb;243:173–180.

Yang H, Pinello CE, Luo J, Li D, Wang Y, Zhao LY, et al. 2013. Small-molecule inhibitors of acetyltransferase p300 identified by high-throughput screening are potent anticancer agents. Mol Cancer Ther. May;12:610–620. Epub 2013/04/30.

Yue M, Jiang J, Gao P, Liu H, Qing G. 2017. Oncogenic MYC Activates a Feedforward Regulatory Loop Promoting Essential Amino Acid Metabolism and Tumorigenesis. Cell Rep. Dec 26;21:3819–3832.

Zhang H, Zha X, Tan Y, Hornbeck PV, Mastrangelo AJ, Alessi DR, et al. 2002. Phosphoprotein analysis using antibodies broadly reactive against phosphorylated motifs. J Biol Chem. Oct 18;277:39379–39387.

