## Supplemental Information for "Repurposing Tranexamic Acid as an Anticancer Agent"

**Supplementary Material**

Table of Contents | Pages

Supplemental Data

Supplemental figure (S) 1:
TA inhibits DNA synthesis without affecting the levels of Cyclin B and Cyclin D1 or the phosphorylation of Akt, ERK and RB 2

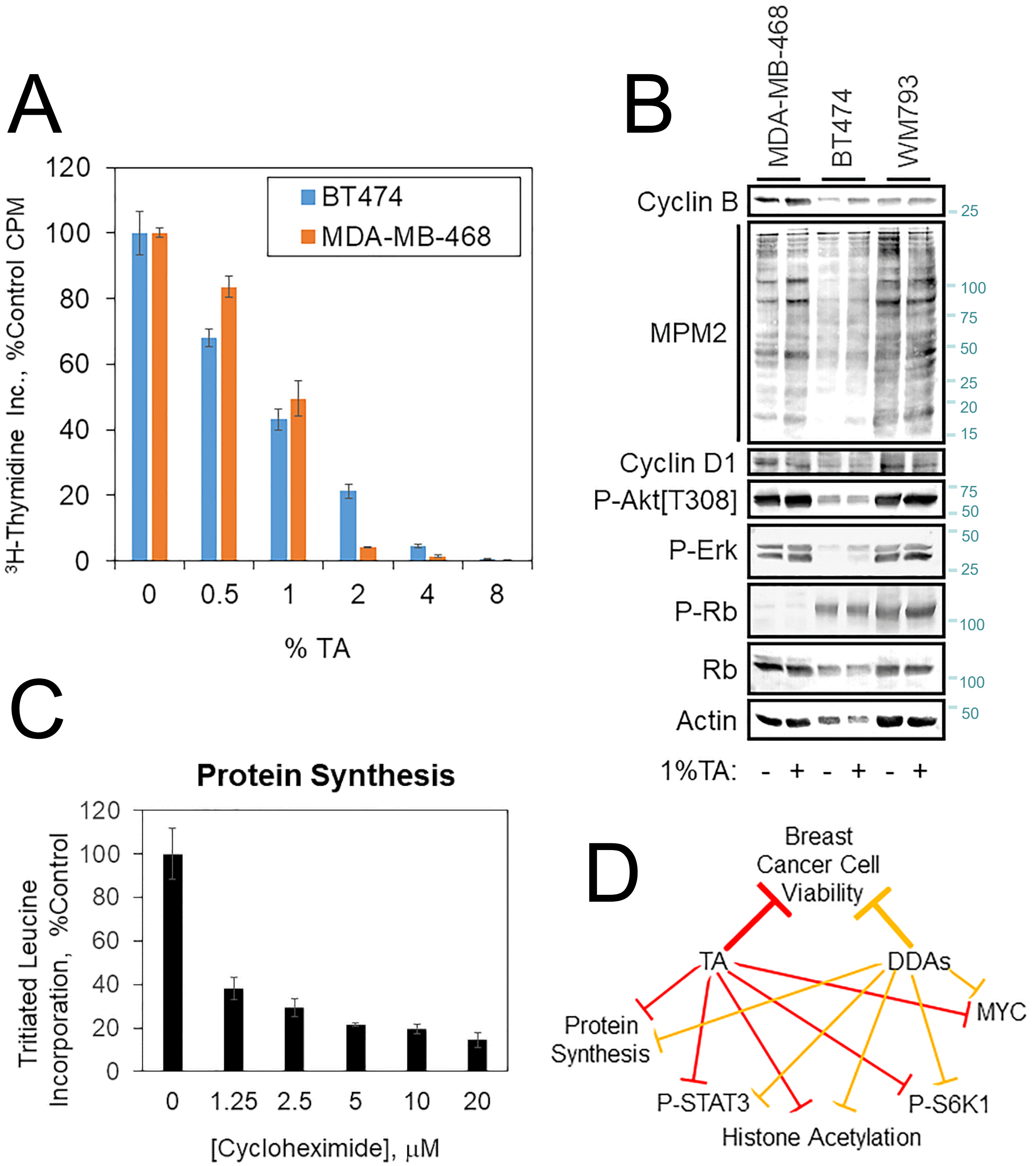


Figure S1: TA inhibits DNA synthesis without affecting the levels of Cyclin B and Cyclin D1 or the phosphorylation of Akt, ERK and RB. A. MDA-MB-468 and BT474 cells were treated for 24 h with the indicated concentrations of TA and the incorporation of tritiated thymidine into DNA was evaluated. B. The indicated cell lines were treated for 24 h with or without 1% TA and cell extracts were analyzed by immunoblot with the indicated antibodies. C. MDA-MB-468 cells were treated for 24 h as indicated and protein synthesis was assayed by tritiated Leu incorporation into protein. Note that 1.25 □M CHX reduces overall protein synthesis by approximately 60%. D. Model displaying the overlapping biochemical mechanisms by which TA and DDAs may suppress breast tumor growth.
